# Superior colliculus modulates cortical coding of somatosensory information

**DOI:** 10.1101/715847

**Authors:** Saba Gharaei, Suraj Honnuraiah, Ehsan Arabzadeh, Greg J Stuart

**Affiliations:** Eccles Institute of Neuroscience, John Curtin School of Medical Research, The Australian National University, Canberra, Australia; Australian Research Council Centre of Excellence for Integrative Brain Function, The Australian National University Node, Canberra, Australia

**Keywords:** Superior colliculus, cortex, sensory processing, optogenetics, thalamus

## Abstract

The cortex sends a direct projection to the superior colliculus. What is largely unknown is whether (and if so how) the superior colliculus modulates activity in the cortex. Here, we directly investigate this issue, showing that optogenetic activation of superior colliculus changes the input-output relationship of neurons in somatosensory cortex during whisker movement, enhancing responses to low amplitude whisker deflections. While there is no direct pathway from superior colliculus to somatosensory cortex, we found that activation of superior colliculus drives spiking in the posterior medial (POm) nucleus of the thalamus via a powerful monosynaptic pathway. Furthermore, POm neurons receiving input from superior colliculus provide excitatory input to somatosensory cortex. Silencing POm abolished the capacity of superior colliculus to modulate cortical whisker responses. Our findings indicate that the superior colliculus, which plays a key role in attention, modulates sensory processing in somatosensory cortex via a powerful disynaptic pathway through the thalamus.

## Introduction

The ability of an organism to attend to, and orient towards, stimuli in the environment is critical for survival. A principal neural substrate for attentional orienting movements is the midbrain structure called the superior colliculus (SC), which receives inputs from multiple sensory modalities and plays an important role in moving the eyes, head and body towards or away from biologically significant stimuli (McHaffie & Stein, 1982; Sparks, 1999; Rowland *et al*., 2007; Gharaei *et al*., 2018). As evidence of its importance, the anatomical structure and input/output architecture of the SC is conserved across a range of mammalian species (May, 2006).

It is well established that SC receives direct input from the primary sensory cortices (Welker *et al*., 1988; May, 2006; Cohen *et al*., 2008; Triplett *et al*., 2012; Castro-Alamancos & Favero, 2016; Zingg *et al*., 2017). What is less clear is whether SC in turn modulates information processing in the cortex. Work in monkeys indicates that SC can modulate activity in higher order cortical areas, with visual responses in the middle temporal area (MT) of monkeys disappearing when lesions of primary visual cortex are combined with lesions of SC (Rodman *et al*., 1990). In contrast, visual responses in the lateral suprasylvian area in cats, which is thought to be analogous to MT in monkeys, are increased by lesions of SC (Smith & Spear 1979; although see Ogino & Ohtsuka 2000). Later work showed that a functional pathway exists from SC to MT through the pulvinar in primates (Berman & Wurtz 2011; although see Stepniewska et al. 1999).

Similar to these earlier studies in primates and cats, more recent work in mice indicates that SC modulates visual responses in higher order cortical visual areas (Tohmi *et al*., 2014) as well as in the postrhinal cortex (Beltramo & Scanziani, 2019). Finally, it has recently been shown that SC can also modulate responses in primary visual cortex in mice, through the dorsolateral geniculate nucleus rather than the pulvinar (Ahmadlou *et al*., 2018). Together, these studies allude to the importance of SC for visual processing in both primary and higher order visual areas in the cortex. What is not known is whether SC also modulates cortical processing from other sensory modalities.

Rodents heavily rely on their whiskers (or vibrissae) to explore and navigate the environment. Sensory information from the whiskers is processed by the whisker associated area of the somatosensory cortex, known as the primary vibrissal somatosensory cortex (vS1; Brecht 2007; Diamond & Arabzadeh 2013). In addition, intermediate and deeper layers of SC also receive sensory information from the whiskers, via a projection from vS1 as well as directly via the trigeminal nucleus of the brainstem (Wise & Jones, 1977; Killackey & Erzurumlu, 1981; Bosman *et al*., 2011; Triplett *et al*., 2012; Castro-Alamancos & Favero, 2016). While several studies have shown that SC neurons are directly activated by whisker deflections (Dräger & Hubel, 1976; Castro-Alamancos & Favero, 2016), it is not known whether activation of SC modulates coding of whisker input in vS1. This issue is the focus of the current study.

## Results

To determine if activation of SC can impact on sensory coding in somatosensory cortex, we expressed ChR2 in intermediate/deep layers of mouse SC. We then performed whole-cell and extracellular recordings to characterize how sensory responses in vS1 cortex were affected by optogenetic activation of SC.

### Activation of SC modulates vS1

To verify that neurons in SC could be reliably driven by light, extracellular recordings were made from SC neurons *in vivo* using multi-electrode optrodes during whisker stimulation (Fig. 1a). Receiver operating characteristic (ROC) analysis indicated that the vast majority of neurons in SC (91%; 50 out of 55) significantly increased their action potential firing in response to brief (15 ms) optogenetic activation (Fig. 1b,c; p < 0.05). Direct activation of SC neurons by light was also verified using whole-cell recordings from SC neurons *in vitro* (Fig. 1d; n=3). To determine if optogenetically activated SC neurons were responsive to whisker input, we identified whisker responsive neurons located in intermediate layers of SC (1.3-2.5 mm from the surface of the brain) using whisker pad vibrations of different amplitudes (Fig. 1e; red). Neurons in SC have large whisker receptive fields and are known to respond robustly to multi-whisker movements (Hemelt & Keller, 2008; Cohen *et al*., 2008). ROC analysis indicated that the majority (80%; 40 out of 50) of SC neurons that responded significantly to light also responded to whisker stimulation (Fig. 1e; green). This led to an upward shift of the input-output relationship of SC neurons to whisker stimuli of different amplitude (Fig. 1f; n=40). Together, these experiments indicate that SC neurons processing sensory input from the whiskers can be reliably activated using optogenetics.

To investigate how activation of SC impacts on sensory coding in somatosensory cortex, whole-cell, loose-patch and extracellular array recordings were made from vS1 while simultaneously activating intermediate/deep layers of SC optogenetically via an optic fiber (Fig. 2a). Brief optogenetic activation of intermediate/deep layers of SC (15 ms) caused increased action potential firing in vS1 neurons (Fig. 2b). Increases in action potential firing in vS1 were observed following optogenetic activation of SC with all three recording techniques (Fig. 2c; extracellular array n=41; loose-patch n=85; whole-cell n=23; p < 0.05). ROC analysis of responses obtained across multiple trials indicated that approximately 60% of vS1 neurons (87 out of 149) showed a statistically significant increase in action potential firing following activation of SC (Supplementary Fig. 1a,b). The median spike latency of vS1 neurons to optogenetic activation of SC was 43.9 ± 9 ms (n=72; see Methods). Increases in action potential firing during optogenetic activation of SC were seen across all cortical depths (Fig. 2d).

**Fig. 1:**
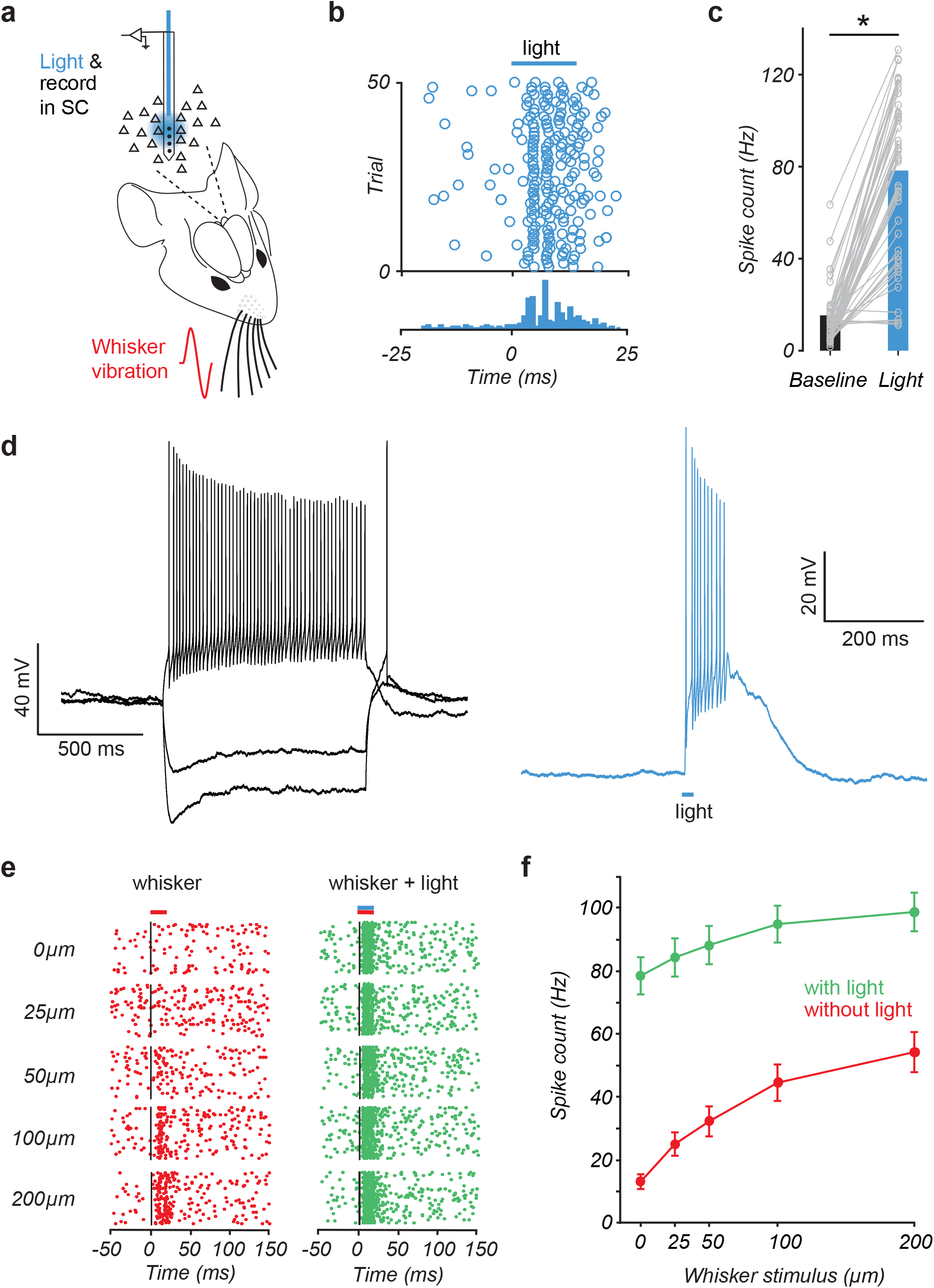
Neurons in SC can be reliably activated by light and whisker stimulation. **a.** Schematic of the experimental arrangement. Extracellular recordings were made from SC during whisker vibration and/or optogenetic activation with light. **b.** Raster plot (top) and peri-stimulus time histogram (bottom) during extracellular recording from a neuron in the intermediate layer of SC using an optrode (2289 μm from the surface of the brain) showing increased spiking in response to light. c. Spiking activity of SC neurons (n=55) increases significantly during light activation. **d.** Left: Voltage responses of a ChR2 expressing neuron in SC to somatic step current injections (−150, −100 & 150 pA). Right: Action potentials evoked in the same neuron in response to light (15 ms; LED power 0.3 mW). **e.** Raster plot of action potential firing in a SC neuron (2130 μm from the surface of the brain) activated by whisker movement of different amplitude alone (red dots; left) and with light (15 ms; green dots; right). **f.** Pooled data showing the impact of light activation (green; 15 ms) on the whisker input-output relationship of whisker responsive SC neurons (red; n=40). Spiking response of each neuron was normalized to the maximum response to whisker stimulation alone. Error bars represent SEM.

**Fig. 2:**
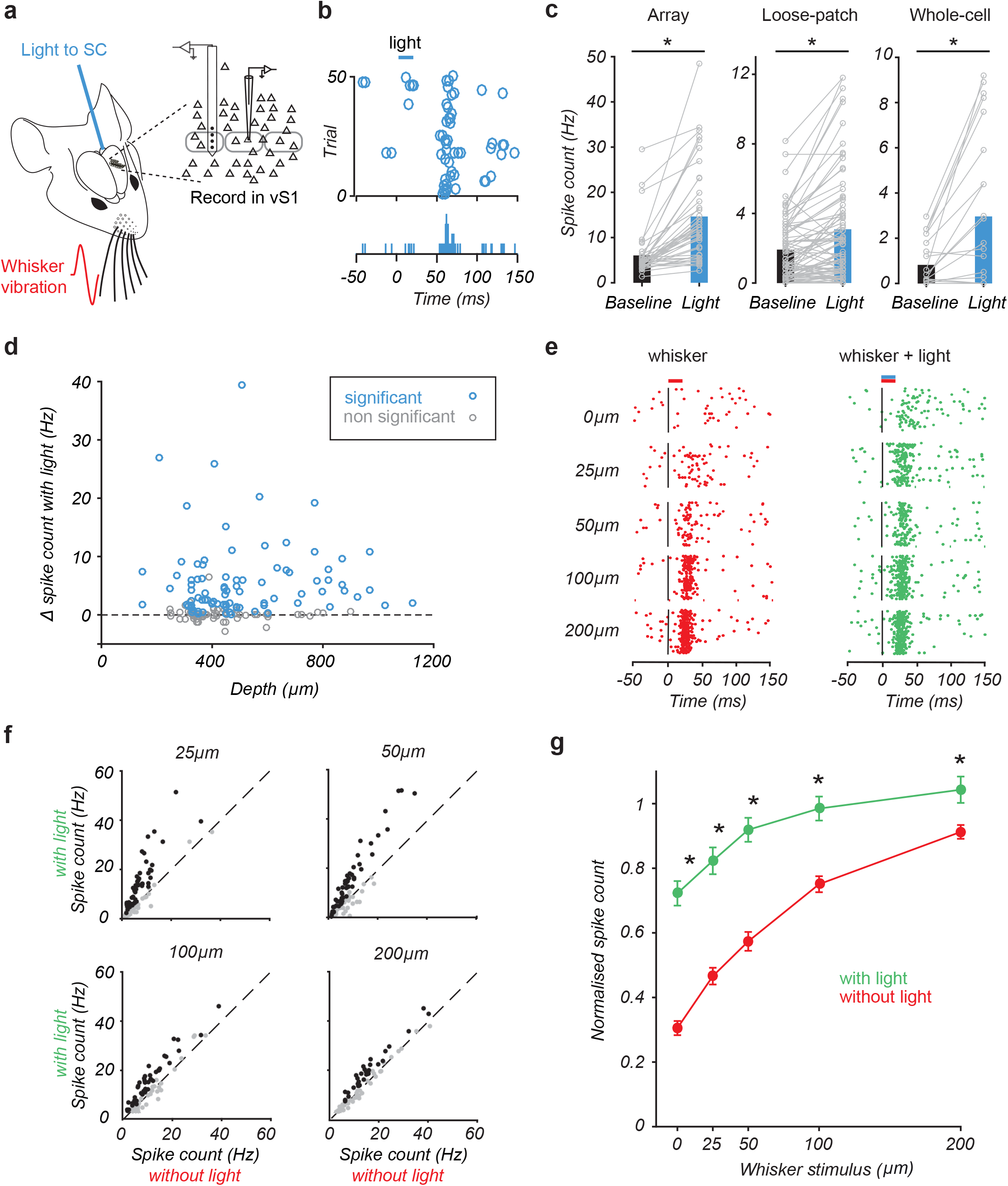
Optogenetic activation of SC modulates vS1. **a.** Schematic of the experimental arrangement. Recordings were made from the vS1 while activating SC optogenetically via an optic fiber in the presence or absence of whisker vibration. **b.** Raster plot (top) and peri-stimulus time histogram (bottom) of a cell in vS1 (513 μm from the surface of the brain) showing increased spiking during SC stimulation (loose-patch configuration). **c.** Spiking of vS1 neurons increases significantly during light activation of SC (extracellular array recordings: n=41; loose-patch recordings: n=85; whole-cell recordings: n=23). **d.** Change in spiking during optogenetic activation of SC compared to baseline versus recording depth in vS1 (n=149). Significant increases in spiking are shown in blue (ROC analyses). **e.** Raster plot of action potential firing in a vS1 neuron (589 μm from the surface of the brain) during whisker movement of different amplitude alone (red dots; left) and with light activation of SC (15 ms; green dots; right). **f.** Plot of action potential firing in whisker responsive vS1 neurons (n=127) with and without light activation of SC during whisker stimulation of different amplitude. Black symbols indicate a significant increase in firing during SC activation (n=101; ROC analysis). **g.** Pooled data showing the impact of SC activation (green; 15 ms) on the whisker input-output relationship (red). Only neurons that were whisker responsive and significantly increase their firing in response to SC activation were included in this analysis (n=101). The spiking of each neuron was normalized to the maximum response to whisker stimulation alone. Error bars represent SEM.

### SC influences processing of somatosensory information in vS1

We next determined how SC activation impacts on responses of vS1 neurons to whisker stimulation. Whole-cell, loose-patch and extracellular array recordings were made from vS1 neurons during whisker vibrations with or without optogenetic activation of SC. In these and all subsequent *in vivo* experiments whisker stimulation (15 ms) was presented simultaneously with optogenetic activation of SC. For each neuron we characterized the spiking response to whisker vibrations of different amplitudes with and without SC activation (Fig. 2e). Activation of SC increased action potential firing during whisker stimulation in 80% of neurons (101 out of 127; ROC analysis) that were whisker responsive (127 out of 149; ROC analysis). Increases in action potential output following SC activation were observed across all intensities of whisker stimulation tested (Fig. 2f). On average, activation of SC caused an upward shift in the input-output relationship of vS1 neurons to whisker stimuli, with the greatest effect observed during low intensity whisker vibrations (Fig. 2g; n=101; p < 0.05). The effect of SC activation on whisker responses was dependent on the whisker input-output relationship, with activation of SC only enhancing whisker responses for whisker vibration amplitudes lower than that evoking the maximal response (Supplementary Fig. 1c). Thus, the greatest impact of SC activation was on whisker defections with the smallest amplitude. Together, these experiments indicate that SC activation leads to an upward shift in the input-output relationship of cortical neurons during somatosensory input, resulting in enhanced responses to low intensity stimuli.

### Circuitry underlying the influence of SC on vS1

We next investigated the circuitry underlying modulation of vS1 by SC. As illustrated schematically in Fig. 3a, there are two main circuits through which SC could impact on sensory processing in vS1 (McHaffie & Stein, 1982; Roger & Cadusseau, 1984; May, 2006; Hemelt & Keller, 2008; Castro-Alamancos & Keller, 2011; Triplett *et al*., 2012). The projection from SC to the facial motor nucleus could lead to whisker movement and thereby modulate vS1 through the conventional ascending pathways via the thalamic ventral posteromedial nucleus (VPM) and posteromedial complex (POm) (Fig. 3a; orange arrows). Alternatively, SC could modulate vS1 through a more direct pathway via the POm (Fig. 3a; purple arrows).

**Fig. 3:**
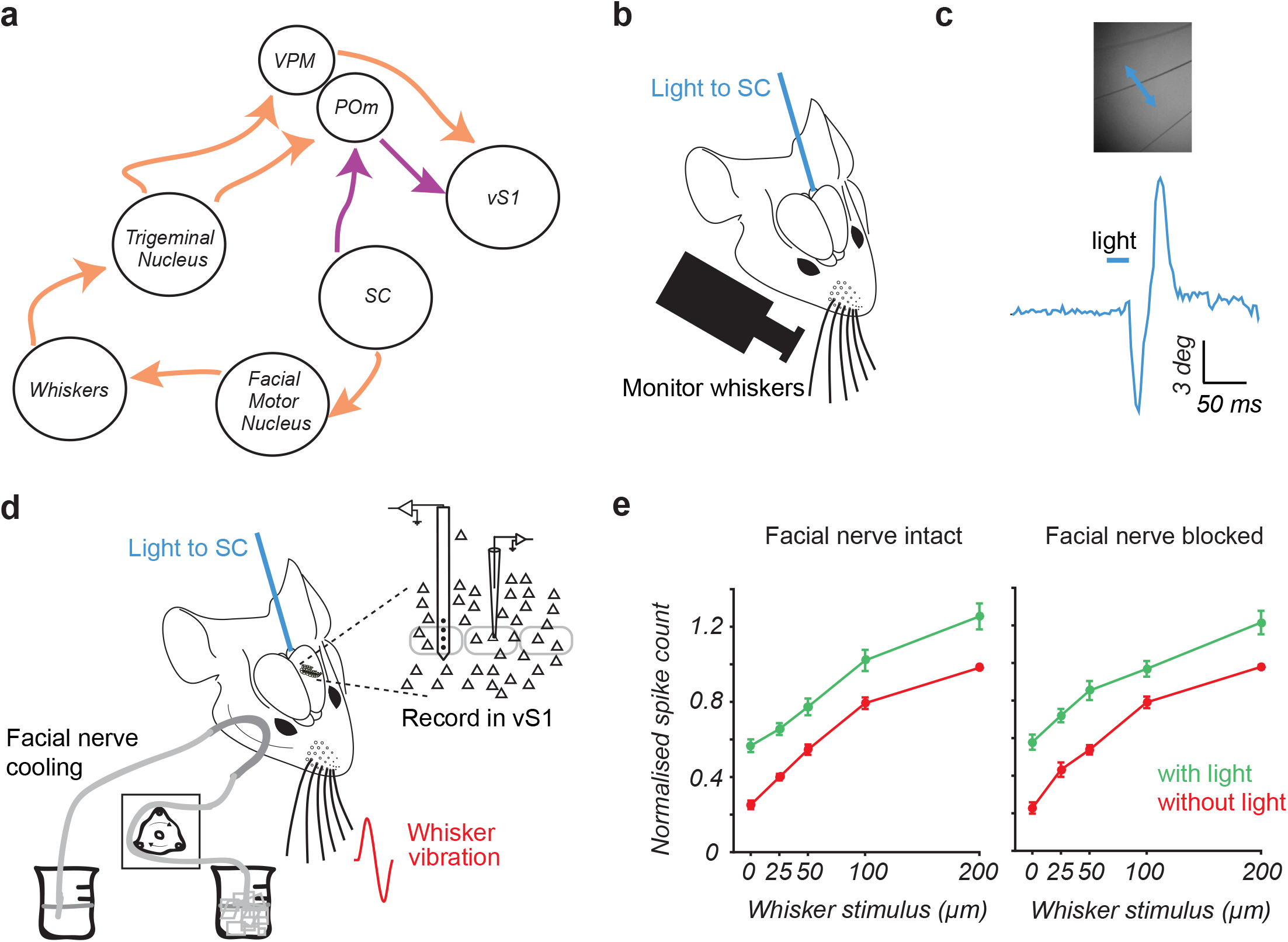
Modulation of vS1 by SC is not due to whisker movement. **a.** Circuits of interest showing two main pathways through which SC could impact on responses in vS1. Modified from Castro-Alamancos and Keller (2011). **b.** Schematic of the experimental arrangement using a high-speed camera to monitor whisker movement while activating SC optogenetically via an optic fiber. **c.** Whisker (top) movement during optogenetic activation of SC (bottom). **d.** Schematic of the experimental arrangement. The impact of optogenetic activation of SC on vS1 responses was measured before and after facial nerve cooling to block whisker movement. **e.** Pooled data showing the impact of SC activation (green; 15 ms) on the whisker input-output relationship of the same whisker responsive vS1 neurons before and after facial nerve inactivation (red; facial nerve cooling: n=16; facial nerve cut: n=8). The spiking of each neuron was normalized to the maximum response to whisker stimulation alone. Error bars represent SEM.

Previous work indicates that micro-stimulation of SC leads to whisker movement (Hemelt & Keller 2008). Consistent with this, optogenetic activation of SC produced whisker movements of short latency (Fig. 3b,c; average onset latency: 22.3 ± 0.22 ms; n=2), suggesting that activation of SC could impact on sensory coding in vS1 by causing whisker movement. We therefore investigated whether blocking activity in the facial nerve on the same side as whisker stimulation impacted on the capacity of SC to modulate sensory coding in contralateral vS1. In rodents, motor commands driving whisker movement arise from the facial motor nucleus and project to the whiskers via the facial nerve, whereas sensory information from the whiskers is conveyed to the trigeminal nucleus via the trigeminal nerve (Semba & Egger, 1986; Hemelt & Keller, 2008; Sachidhanandam *et al*., 2013; Heaton *et al*., 2014; Kaneshige *et al*., 2018). As a result, it is possible to abolish whisker movements by blocking facial nerve activity while maintaining sensory input to vS1 from the whiskers. Whisker movements following optogenetic activation of SC were abolished by cutting or reversible cooling the facial nerve (Fig. 3d). Importantly, this procedure had no impact on the capacity of optogenetic activation of SC to modulate the whisker input-output relationship of neurons in vS1 (Fig. 3e; n=8 facial nerve cut; n=16 facial nerve cooling). These data indicate that SC does not modulate vS1 through generation of whisker movement. This finding is perhaps not surprising given that whisker protractions induced by SC activation are of relatively low velocity (Hemelt & Keller, 2008), and therefore not expected to have a significant impact on spiking in vS1 neurons (Arabzadeh *et al*., 2005).

We next investigated the possibility that SC modulates vS1 through an indirect pathway via POm (Roger & Cadusseau, 1984; Castro-Alamancos & Keller, 2011; Stein, 2012). Consistent with this idea, optogenetic activation of SC led to increased firing of POm neurons (Fig. 4a-c; n=38; p < 0.05). Across the population of POm neurons recorded using both loose-patch and array recording, ROC analysis indicated that optogenetic activation of SC increased action potential firing in 66% of POm neurons (Fig. 4c; 25 out of 38). The facial nerve on both sides of the snout were cut in these experiments, ruling out the possibility that activity in POm was driven by whisker movement. The median spike latency of POm neurons following SC activation was 23 ± 4.0 ms (n=25). For each neuron in POm, we established a stimulus response function to whisker vibrations of different amplitude in the presence and absence of optogenetic activation of SC (Fig. 4d). SC activation increased action potential firing to whisker stimulation in almost all POm whisker responsive neurons (15 out of 16; ROC analysis). These experiments indicate that, as seen in vS1, SC activation causes an upward shift in the input-output relationship of POm neurons to whisker input (Fig. 4e; n=15). In summary, these experiments provide functional evidence that POm neurons are reliably driven by optogenetic activation of the SC.

**Fig. 4:**
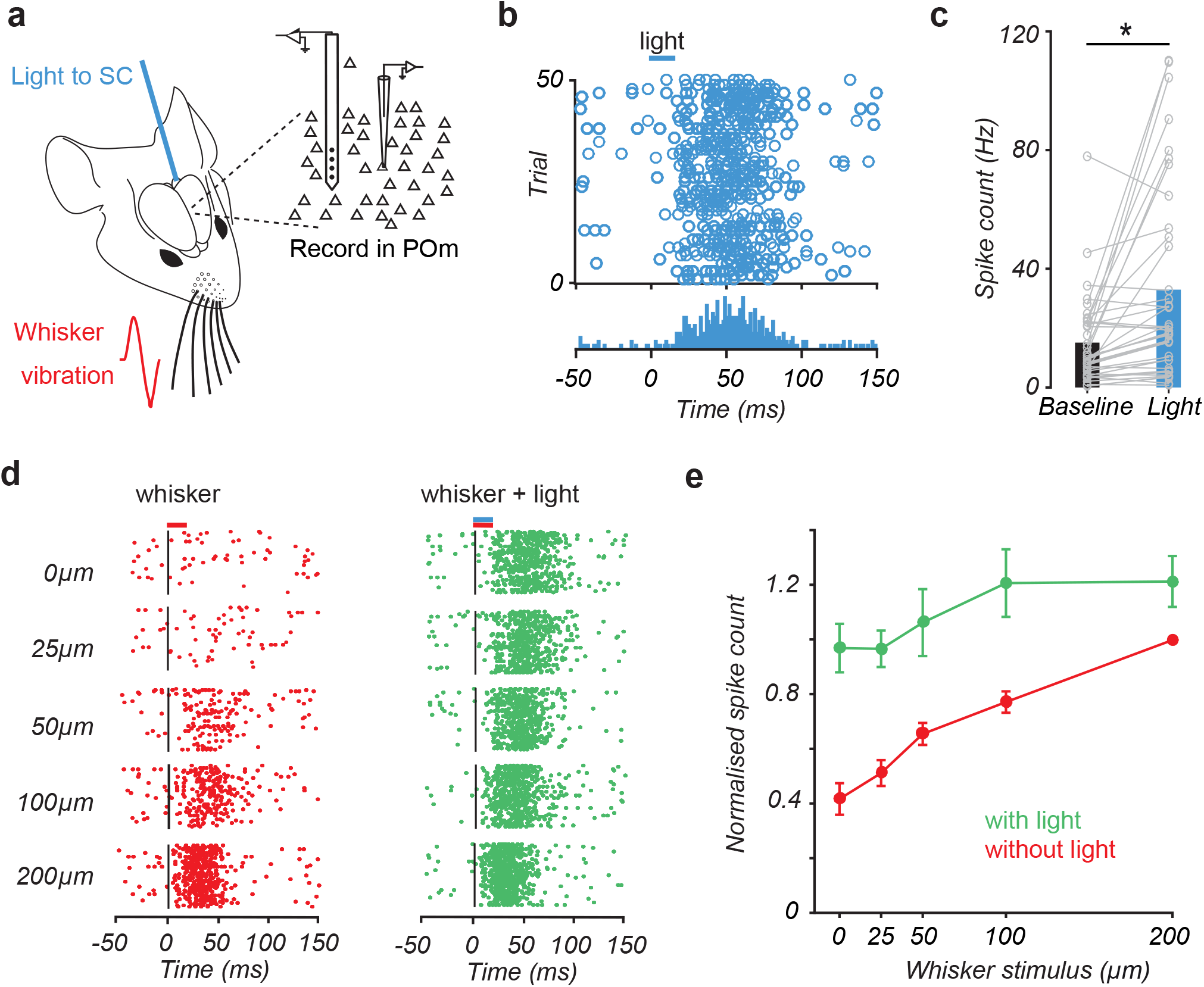
Activation of SC drives activity in POm. **a.** Schematic of the experimental arrangement. After cutting the facial nerves, recordings were made from the POm while activating SC optogenetically through an optic fiber in the presence or absence of whisker vibration. **b.** Raster plot (top) and peri-stimulus time histogram (bottom) of an extracellular array recording from a POm neuron (2407 μm from the surface of the brain) showing increased action potential firing in response to SC activation. **c.** Spiking of POm neurons (n=38) increases significantly during light activation of SC. **d.** Raster plot of action potential firing in a POm neuron (same neuron as panel B) during whisker movement of different amplitude alone (red dots; left) and with light activation of SC (green dots; right). **e.** Pooled data showing the impact of SC activation (green; 15 ms) on the whisker input-output relationship of whisker responsive POm neurons (red; n=15). Spiking of each neuron was normalized to the maximum response to whisker stimulation alone. Error bars represent SEM.

### SC sends a direct projection to POm

We next investigated if SC sends a direct, monosynaptic projection to neurons in POm using whole-cell recordings from POm neurons *in vitro* (Fig. 5a). Brief (2 ms) optogenetic activation of SC axons evoked excitatory postsynaptic potentials (EPSPs) in 73% of POm neurons (Fig. 5b, left; 16 out of 22 neurons). These neurons all received monosynaptic input from SC, as EPSPs remained in the presence of TTX plus 4-AP (Fig. 5b, right; n=16). No polysynaptically driven cells in POm were observed. The properties of POm neurons receiving input from SC were similar to those of cells that did not receive input from SC (Supplementary Fig. 2). Of the cells receiving monosynaptic SC input, the majority (69%; 11 out of 16) responded to SC input by generating action potentials in response to the lowest LED light intensity tested (LED power: 0.8 mW), whereas the remaining cells (5 out of 16) evoked graded responses, with an average amplitude of 2.2 ± 0.6 mV (LED power: 0.8 mW). We therefore classified POm cells into two groups: Cells that generated suprathreshold spiking in response to low intensity LED stimulation were classified as receiving “strong” SC input, whereas cells that generated small, subthreshold EPSPs in response to low intensity LED stimulation were classified as receiving “weak” SC input. The passive and active properties of POm cells receiving strong and weak input from SC were found to be similar, suggesting they do not represent different neuronal cell types (Supplementary Fig. 3). When tested with very low LED intensities (less than 0.4 mW) POm neurons receiving strong input from the SC generated graded changes in EPSP amplitude (Fig. 5c, left), but with a very different dependence on LED power compared to cells receiving weak SC input (Fig. 5d). To investigate whether SC also projects to VPM, we made recordings from neurons in VPM (Fig. 5e). No excitatory synaptic responses were observed in any VPM neurons using the highest LED intensity available (5 mW; n=5). Figure 5f summarizes responses obtained from POm and VPM during activation of SC at the highest LED power tested (5 mW). Together, these data indicated that SC powerfully drives the majority of POm neurons via a direct monosynaptic projection, but does not activation neurons in VPM.

**Fig. 5:**
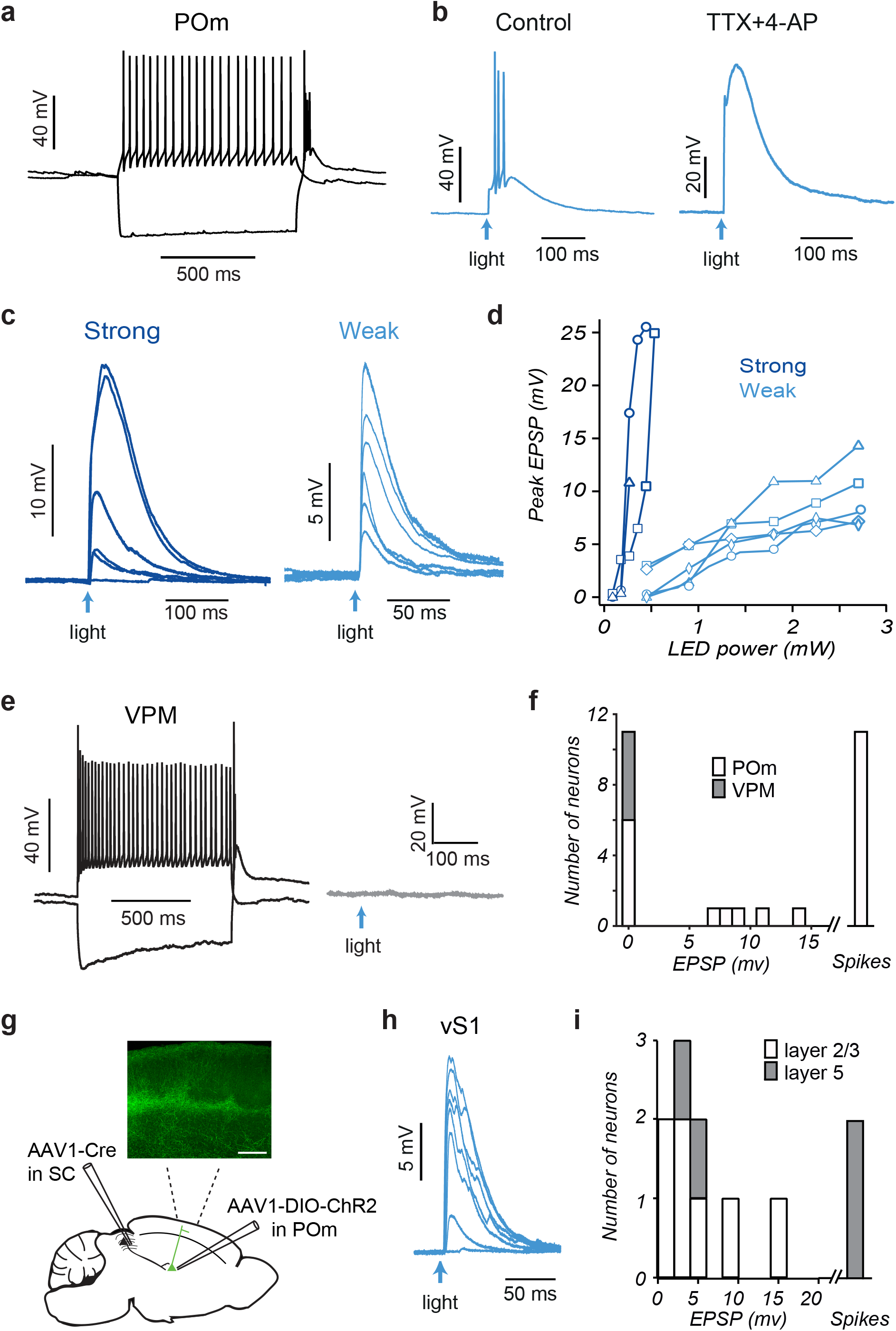
POm, but not VPM, receives direct, monosynaptic input from SC. **a.** Response of a POm neuron to somatic depolarizing (+300 pA) and hyperpolarizing (−400 pA) current steps. **b.** Synaptic response from a POm neuron to photo-activation of SC axons (arrow; 2 ms; 0.8 mW) in control and in the presence of TTX and 4-AP. **c.** Graded synaptic responses in POm neurons receiving “strong” (left; LED power 0.1 to 0.42 mW) and weak (right; LED power 0.42 to 2.72 mW) input from SC. **d.** Summary of EPSP amplitudes plotted as a function of LED power in POm neurons receiving “strong” (dark blue) and “weak” (light blue) SC input. **e.** Right: Response of a VPM neuron to somatic depolarizing (+300 pA) and hyperpolarizing (−400 pA) current injection. Left: Voltage responses of a VPM neuron to photo-activation of SC axons at the maximum LED power (2 ms; 5 mW). **f.** Histogram of EPSP amplitude or spiking in VPM (n=5) and POm (n=22) neurons during optogenetic activation of SC axons at the highest LED power tested (5 mW). **g.** Schematic of the experimental paradigm. Expression of Cre recombinase (AAV1.hSyn.Cre.WPRE.hGH) in SC was coupled with expression of Cre-dependent ChR2 (AAV1-Ef1a-DIO-hChR2(E123A)-EYFP) in POm. The inset shows EYFP expressing POm axons in vS1. **h.** Synaptic responses in a layer 2/3 vS1 neuron to optogenetic activation of POm axons (2 ms; 0.72 to 5 mW). **i.** Histogram of EPSP amplitude or spiking in layer 2/3 or layer 5 vS1 neurons during optogenetic activation of the axons of POm neurons receiving direct input from SC (2 ms; 5 mW).

We next investigated whether SC is di-synaptically connected to vS1 through POm. To investigate this, we used an AAV-mediated anterograde trans-synaptic tagging method (Zingg *et al*., 2017). Cre-dependent expression of ChR2 in POm was driven by anterograde trans-synaptic transfection of Cre recombinase in SC (Fig. 5g). Histological analyses revealed cell bodies expressing EYFP fluorescence in POm with light activation of these neurons leading to action potential generation (n=4). EYFP fluorescent axons were also found in vS1 (Fig. 5g, insert). Whole-cell recordings from neurons in vS1 *in vitro* were used to determine if these POm axons provide functional input to vS1. Brief (2 ms) optogenetic activation of the axons of POm neurons receiving direct input from the SC evoked EPSPs in 71% of layer 2/3 (5 out of 7) and 100% of layer 5 neurons (4 out of 4) in vS1 (Fig. 5h,i). In summary, these experiments confirm that POm neurons receiving direct, monosynaptic SC input project to multiple layers in vS1.

### Silencing POm abolishes SC responses in vS1

We next tested if activation of POm is required for SC modulation of vS1. To investigate this we silenced POm by pressure injection of a small volume (100-200 nl) of lidocaine into POm while recording the impact of optogenetic activation of SC on responses in vS1 (Fig. 6a). The facial nerve on both sides of the snout was cut in these experiments. Figure 6b (left) shows extracellular spiking activity of a representative vS1 neuron to optogenetic stimulation of SC. Light responses in this vS1 neuron were abolished after lidocaine was injected into POm (Fig. 6b, right). Pooled data indicated that inactivation of POm lead to a statistically significant reduction in the response of vS1 neurons to optogenetic activation of SC (Fig. 6c; n=36; p < 0.05). We next tested the impact of inactivation of POm on vS1 responses during brief deflections of the whiskers. SC activation increased action potential firing to whisker stimulation in 79% of vS1 whisker responsive neurons in these experiments (23 out of 29; ROC analysis). Silencing POm essentially abolished the impact of SC activation on vS1 neurons during whisker stimulation (Fig. 6d, green; n=23; p < 0.05). Importantly, inactivation of POm had no impact on baseline activity of vS1 neurons in these experiments (Fig. 6d, red; n=23; p > 0.05). Together, these results provide direct evidence that SC modulates vS1 via an indirect pathway through POm of thalamus.

**Fig. 6:**
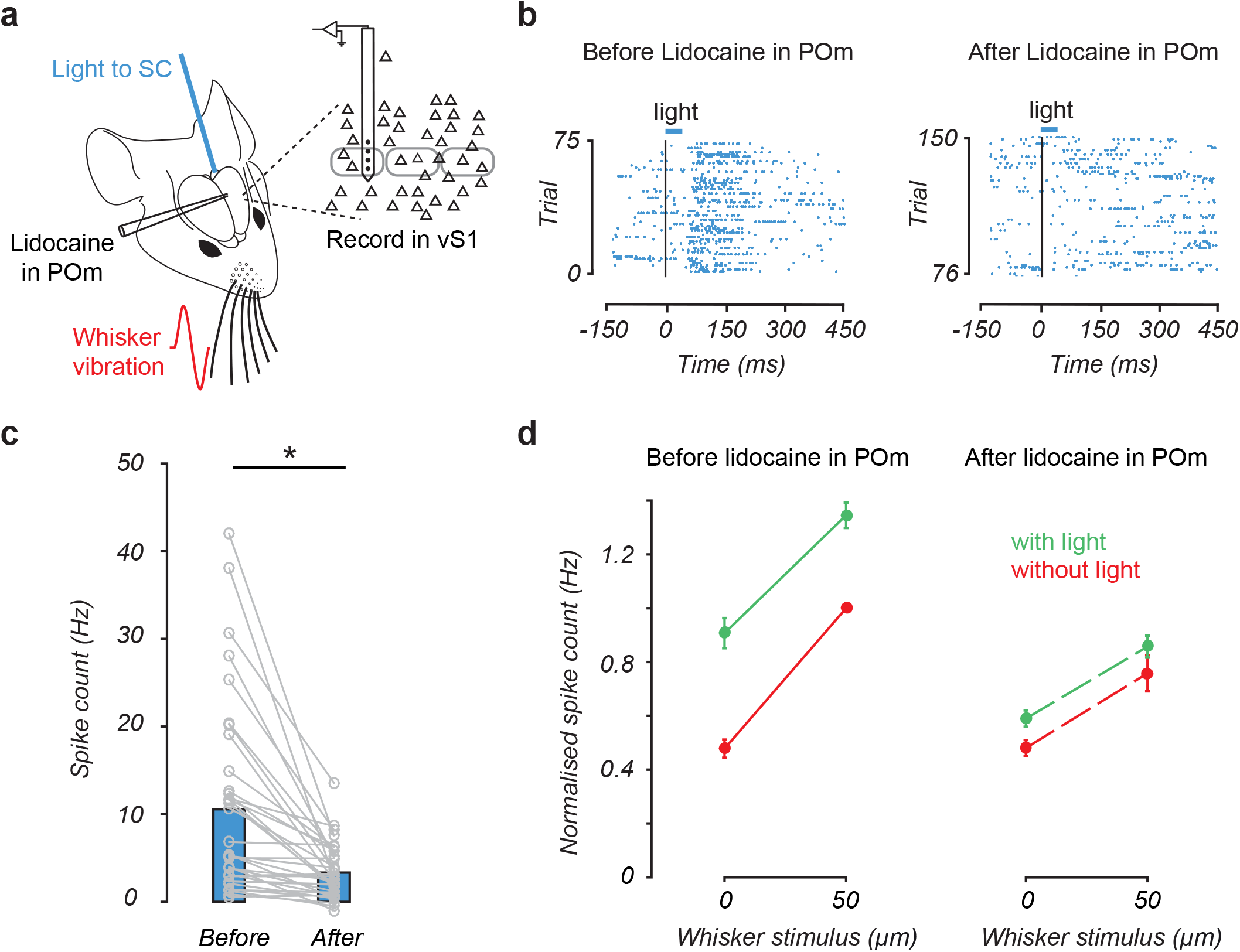
Inactivation of POm abolishes the impact of SC on vS1 neurons. **a.** Schematic of the experimental arrangement. After cutting the facial nerve, recordings were made from vS1 while activating SC optogenetically through an optic fiber in the presence or absence of whisker vibration. POm was silenced by lidocaine injection. **b.** Raster plot during extracellular array recording of spiking in a vS1 neuron (787 μm from the surface of the brain) in response to optogenetic activation of SC in control (left) and following inactivation of POm via lidocaine injection (right). **c.** Plot of baseline-subtracted responses to SC activation before and after lidocaine injection in POm (n=36). **d.** Pooled data (n=23) on the impact of SC activation (green) on spiking in whisker responsive vS1 neurons during small amplitude whisker movement (red) in control (left) and following inactivation of POm via lidocaine injection (right). The spiking activity of each neuron was normalized to the maximum response to the whisker stimulation alone in control. Error bars represent SEM.

## Discussion

Here we directly test the impact of SC on cortical function. While it has long been known that the cortex projects to SC, what is only now becoming clear is that SC also modulates activity in the cortex. A number of recent studies have identified the capacity of SC to influence cortical processing of visual input (Beltramo & Scanziani 2019; Ahmadlou et al. 2018). By combining optogenetic activation of SC with recordings in vS1 of mice, we show here that SC also modulates cortical processing of somatosensory input. This effect of SC on responses in vS1 was not a result of SC driving whisker movement, but instead was mediated via an indirect pathway through POm of the thalamus.

It is well-established that SC is involved in orienting behaviors and directing attention to relevant sensory information (Krauzlis *et al*., 2013), with early work by Schneider (1969) suggesting that in rodents SC is involved in the spatial localization of a stimulus. The multisensory nature of SC makes it an ideal structure for orienting responses towards or away from a salient stimulus by initiating movement of the eyes, whiskers, head and body. Indeed, our observation that optogenetic activation of SC in mice leads to whisker movement is consistent with earlier work showing that micro-stimulation of SC generates whisker and saccadic eye movements in rats (McHaffie & Stein, 1982; Hemelt & Keller, 2008). Sustained protractions evoked by SC stimulation are different from rhythmic protractions caused by stimulation of motor cortex (Hemelt & Keller, 2008). These SC evoked movements were shown to be independent of the motor cortex and increased in magnitude with increasing stimulation intensity (Hemelt & Keller, 2008). These data suggest that the function of whisker movements caused by SC may be to position them relative to an object that has attracted the animal’s attention or in anticipation of head movements rather than sensory coding (Barth & Schallert, 1987; Towal & Hartmann, 2006; Hemelt & Keller, 2008).

Despite the importance of the SC in evoking whisker movement, our experiments show that inactivating the facial nerve, and thereby blocking whisker movement, had no impact on SC-evoked responses in vS1. Instead, we show that activation of SC drives activity in POm *in vivo* (Fig. 4) via a direct, monosynaptic pathway (Fig. 5). Furthermore, we show that inactivation of SC essentially abolishes the capacity of SC to modulate responses in vS1 (Fig. 6). These data indicate that SC modulates vS1 via an indirect pathway through POm of thalamus. This observation is consistent with an earlier anatomical study that identified projections from SC to the rostral sector of POm (Roger & Cadusseau, 1984).

It is well established that POm not only sends a direct input to vS1, but also modulates sensory processing in vS1 (Viaene *et al*., 2011; Castejon *et al*., 2016; Mease *et al*., 2016; Zhang & Bruno, 2019). Consistent with these studies, activation of SC increases activity of vS1 neurons, leading to larger whisker evoked responses particularly during small whisker movements. This effect of SC would be expected to enhance the capacity of the cortex to detect weak sensory input and may therefore play a role in directing attention to relevant stimuli. Furthermore, by modulating coding in vS1, SC may also play a role in feature detection.

Using whole-cell recordings *in vitro,* we show that SC is di-synaptically connected to vS1 through POm. We show that SC sends a monosynaptic projection to neurons in POm, but not VPM, and that POm neurons receiving direct, monosynaptic input from SC project to vS1. Interestingly, we observed three groups of cells in POm: One group received strong SC input, another weak SC input, with a third group that did not receive input from SC. These different populations of POm neurons had similar active and passive properties, suggesting they are likely to be of the same cell type. Further experiments will be required to determine the functional role of these different POm populations. Interestingly, optogenetic activation of SC axons did not lead to polysynaptic responses in POm neurons that did not receive monosynaptic SC input, despite the fact that the majority of cells generated action potentials in response to SC input. This finding is consistent with earlier work indicating that POm neurons have very small or no recurrent connectivity with each other (Deschênes *et al*., 1998; Ohno *et al*., 2012).

Thalamus routes sensory signals to the cortex and thus sits in a strategic position to modulate cortical state and the efficiency of sensory information processing (Sabri & Arabzadeh, 2018). Compared to VPM, POm receives weaker and more diffuse sensory projections from the trigeminal nucleus and projects mainly to layer 1, layer 5a and inter-barrel regions of layer 4 in vS1 (Kichula & Huntley, 2008; Furuta *et al*., 2009; Bosman *et al*., 2011). These observations indicate that POm axons are not distributed evenly across all cortical layers, yet we found that the impact of SC activation on spiking in vS1 was not dependent on recording depth (Fig. 2d). Similarly, we found that the axons of POm neurons receiving direct, monosynaptic input from SC provided excitatory input to neurons in both layer 2/3 and layer 5 (Fig. 5i). These finding are in line with earlier work showing that micro-stimulation of POm evokes EPSPs in neurons located in all cortical layers of vS1 (Viaene *et al*., 2011).

Our observation that whisker-evoked responses in vS1 neurons are enhanced by optogenetic activation of SC is consistent with earlier studies investigating the impact of POm on vS1. This earlier work indicates that whisker-evoked responses in vS1 neurons in both mice and rats are amplified by POm activation during whisker stimulation (Mease *et al*., 2016; Zhang & Bruno, 2019). Sensory enhancement caused by POm is accompanied by prolongation of cortical responses over long time periods after whisker stimulation (Mease *et al*., 2016). This prolonged activity in response to POm activation may prime the cortex for a behavioral response and has been shown to be critical for long-term potentiation of whisker inputs (Gambino *et al*., 2014). Together, these studies suggest that the effect of SC activation on vS1 through POm may be to enhance and sustain cortical sensory signals and thereby emphasize and direct attention to salient sensory information. Consistent with this idea, enhanced population activity is observed in vS1 during simple forms of attention such as sensory prioritization (Lee *et al*., 2016) and temporal cueing (Lee *et al*., 2019). These findings are in line with earlier work in primates (Muller *et al*., 2005; Cavanaugh *et al*., 2006; Lovejoy & Krauzlis, 2010; Zénon & Krauzlis, 2012; Herman & Krauzlis, 2017) as well as attentional gain modulation seen in mouse visual cortex and thalamus (Wimmer *et al*., 2015; McBride *et al*., 2019).

The POm in the rodent somatosensory system is thought to be analogous to the pulvinar in the primate visual system (Viaene *et al*., 2011). Although the function of POm is still controversial (Ahissar & Oram, 2015; Mease *et al*., 2016), the role of the pulvinar in perception, selective visual attention and visual saliency is better understood (Robinson & Petersen, 1992; Snow *et al*., 2009; Berman & Wurtz, 2011).

Given these findings in primates, it is perhaps not surprising that SC, which is known to be involved in attention, projects to POm in rodents. POm not only projects to vS1, but also to striatum, secondary somatosensory cortex, as well as other cortical areas, such as motor and auditory cortices (Ohno *et al*., 2012). Our finding that POm receives direct input from SC, suggests that beyond its involvement in somatosensory perception, activity in POm may generate a general priming signal that acts to modulate sensory processing in multiple cortical areas (Mease et al. 2016).

## Online Methods

### Animals

A total of 76 adult male C57BL6/J mice (age between 4 and 6 weeks) were used in this study. Mice were housed in a controlled environment with a 12-hour light-dark cycle and all animal procedures were approved by the Animal Experimentation Ethics Committee of the Australian National University.

### Viral injections

Glass pipettes (Drummond), pulled on a microelectrode puller (Sutter Instrument Co.; P-87, USA) and broken to give a diameter of around 20 μm, were back-filled with mineral oil and front-loaded with viral suspension (AAV1-hSyn.ChR2(H134R)- eYFP.WPRE.hGH, AAV1.hSyn.Cre.WPRE.hGH, AAV1-Ef1a-DIO-hChR2(E123A)-EYFP; University of Pennsylvania, USA). Animals were placed in a chamber to induce light anesthesia via brief exposure to isoflurane (3.5% in oxygen) then mounted in a stereotaxic frame with anesthesia continued using isoflurane (1-1.5% in oxygen) delivered through a nose cone. Throughout surgery mice were placed on a servo-controlled heating blanket (Harvard instruments) to maintain a steady body temperature near 37°C. For expression of ChR2 in SC, a craniotomy with a diameter of 1 mm was performed above the left SC using stereotaxic coordinates (centered at 0.5 mm anterior to lamda and 1.5 mm lateral to the midline) and a small amount (100-180 nl) of viral suspension (AAV1-hSyn.ChR2(H134R)-eYFP.WPRE.hGH) was injected (36.8 nl per minute; Nanoject II; Drummond) into intermediate/deep layers of SC (1.5-2.0 mm from the surface). In experiments where ChR2 was expressed in POm neurons receiving direct input from SC, AAV1.hSyn.Cre.WPRE.hGH was injected into the left SC (200 nl) and two weeks later a craniotomy with a diameter of 1 mm was performed above the left POm (centered at 1.7 mm posterior to bregma and 1.2 mm lateral to the midline) and a small amount (345-690 nl) of Cre-dependent viral suspension (AAV1-Ef1a-DIO-hChR2(E123A)-EYFP) was injected (36.8 nl per minute). Following viral injections the scalp incision was closed and ketoprofen (5 mg/kg; subcutaneous) was given for pain relief. At the completion of surgery mice were returned to their cage and placed on a heating pad to recover.

### Surgery for in vivo experiments

Three to four weeks after viral injection of AAV1-hSyn.ChR2(H134R)-eYFP.WPRE.hGH into SC, anesthesia was initially induced with brief exposure to isoflurane (3.5% in oxygen) and maintained by intraperitoneal administration of urethane (500 mg/kg) together with chlorprothixene (5 mg/kg). The level of anesthesia was regularly monitored by checking hind paw and corneal reflexes, and maintained at a stable level by administering top-up dosages (10% of original) as required. Atropine (0.3 mg/kg, 10% w/v in saline) was administered subcutaneously to reduce secretions. The animal was placed on a servo-controlled heating blanket (Harvard instruments) to maintain a steady body temperature near 37°C. A custom-built head holder was glued to the skull and stabilized with dental cement. The head holder was mounted on a steel frame to minimize head movement. A craniotomy (diameter 1 mm) was performed above the left SC at the same location where viral injections had been performed (0.5 mm anterior to the lambda and 1.5 mm lateral to the midline). Another craniotomy (diameter 2 mm) was performed above the left vS1 (1.5 mm posterior to the bregma and 3 mm lateral to the midline). Saline was applied to exposed areas so they remained moist. In a subset of experiments, a craniotomy was performed above POm (diameter 2 mm, centered 1.7 mm posterior to bregma and 1.2 mm lateral to the midline). Dura mater was left intact for all areas. At the end of the experiment the animal was euthanized with an overdose of sodium pentobarbitone (100 mg/kg intraperitoneal; Lethobarb; Verbac Australia, NSW, AUS). To confirm viral expression and electrode location, animals were perfused trans-cardially with 0.9% sodium chloride solution and then 4% paraformaldehyde (PFA). The brain was removed from the skull and kept in PFA overnight. Coronal slices (100 μm thick) were prepared and examined under a confocal microscope (A1 Nikon).

### Multi-electrode array recording

Extracellular single unit activity was recorded with 4-channel linear Neuronexus silicon electrodes (spacing between electrodes: 100 μm). Recordings from SC were performed using an optrode (a 4-channel linear silicon electrode equipped with an optic fiber; Neuronexus) inserted vertically into SC (1.5-2.5 mm from the surface). During recordings in vS1, Neuronexus silicon electrodes were inserted at an oblique angle of 45° (0.15-1.1 mm depth). Signals from all 4 electrodes of the array were simultaneously amplified, filtered (250-5000 Hz) and were continuously recorded onto disk at a sampling rate of 40 kHz (Plexon amplifier). Data were sorted off-line to identify spiking activity on each channel. A negative threshold of 4 standard deviations of the background noise was used to detect spikes on each channel. For recordings in POm, electrodes were inserted to a depth of 2.4-3.0 mm from the surface of the brain. In a subset of these experiments, to determine the location of recording sites the multi-electrode array was dipped in fluorescent dye (Fast DiI oil; ThermoFisher Scientific) prior to insertion, with electrode location verified post-hoc.

### In vivo whole-cell and loose-patch recording

Whole-cell (Margrie *et al*., 2002) and loose-patch recordings were used to record the subthreshold and spiking activity of vS1 neurons. Patch pipettes were pulled from borosilicate glass and had open tip resistances of 5-7 MΩ when filled with an internal solution containing (in mM): 130 K-gluconate, 10 KCl, 10 HEPES, 4 MgATP, 0.3 Na_2_GTP, 15 Na_2_Phosphocreatine (pH 7.25 with KOH, osmolality ~290 mOsm). Electrodes were inserted into the brain at an oblique angle (30–45°) and lowered rapidly using a Sutter micromanipulator with high positive pressure (~200 mmHg) to pass the dura mater. The pressure was then dropped to 30 mmHg and the pipette advanced at a speed of ~2 μm/s while searching for neurons. Pipette resistance was constantly monitored in voltage clamp by applying 10 mV voltage pulses with a duration of 20 ms at a frequency of 25 Hz. For loose-patch (juxta-cellular) recordings the final seal resistance was more than 40 MΩ. For whole cell recordings, following contact with a cell the command potential was hyperpolarized to −65 mV and constant suction of up to 70 mmHg applied. After a gigaseal was established, brief pulses of suction were applied to the pipette to rupture the membrane. Both loose-patch and whole-cell recordings were performed in current clamp using a BVC-700A amplifier (Dagan Corporation, Minneapolis, MN). Voltage was low-pass filtered at 10 kHz using a Bessel filter prior to being digitized at 40 kHz using either an ITC-18 (Instrutech) or a PCIe-6321 data acquisition board (National Instruments). Current and voltage signals were acquired by a PC computer running Axograph acquisition software (Axograph Scientific, Sydney, Australia).

### In vivo optogenetic light activation of SC

For simultaneous recording and light activation of SC, we used an optrode equipped with an optic fiber (125 μm core diameter; Neuronexus). The optrode optic fiber was connected to a 470 nm LED (Thorlabs), with the power of blue light out of the fiber tip set to 1.8 mW. The LED was controlled through a National Instrument board using programs written in Matlab (MathWorks, Inc., Natick, MA). For activating SC while recording from vS1 or POm, photo activation was delivered through an optic fiber (200 μm core diameter) connected to a 470 nm LED (Thorlabs). In this case, the power of blue light from the tip of the optic fiber was 2.9 mW. Both the optrode and the optic fiber were inserted vertically into the SC (1.5-2.5 mm from the surface). The duration of SC light activation was 15 ms.

### Whisker Stimulation

Brief (15 ms) whisker deflections were applied to the right vibrissal pad (contralateral to the recording site) using a light-weight fine mesh plate glued to a piezoelectric ceramic (Morgan Matroc, Bedford, OH). The piezo was driven by an amplifier (PiezoDrive, amplification gain of 20) controlled by commands generated in MATLAB and sent to the analogue output of a PCIe-6321 data acquisition board (National Instruments; 20 kHz sampling rate). The voltage signal had a Gaussian waveform, which produced a brief deflection (6 ms rise, 9 ms decay) with minimal ringing (Ranjbar-Slamloo & Arabzadeh, 2017). Five amplitudes (0, 25, 50, 100 and 200 μm) were delivered in the vertical direction either with or without optogenetic activation of SC (10 stimulus conditions in total). In all *in vivo* experiments whisker stimulation was presented simultaneously with optogenetic activation of SC. Stimuli were applied in a pseudo-random order with 50 repetitions per condition with 850 ms interstimulus interval. Each episode of recording included 500 trials.

### Whisker tracking

For whisker tracking, whiskers were trimmed to the level of the facial hairs except for C row, which was illuminated from below by visible light. High speed videos were captured at 400 frames per second with a CMOS camera (PhotonFocus, Lachen, Switzerland) mounted on a Leica M80 stereomicroscope during a 1 second period (0.5 seconds before and 0.5 seconds after SC activation). Frame acquisition was triggered by a National Instrument board. We used automated software (Clack et al. 2012) to calculate whisker angle and curvature.

### Cutting or reversible inactivation of the facial nerve

In some experiments, mice underwent bilateral facial nerve (VII) transection before performing the craniotomy and recordings. Two incisions were made in the skin covering both cheeks to expose the facial nerves. The facial nerves on both sides were then cut using microsurgical scissors under a dissecting microscope using procedures similar to that described in previous papers (Sachidhanandam *et al*., 2013; Heaton *et al*., 2014). In a subset of these experiments, reversible inactivation of the facial nerve was achieved by nerve cooling. The facial nerve was exposed and a custom-made stainless steel cryoloop was placed over the exposed nerve. The cooling procedure was similar to that described previously (Lomber *et al*., 1999; Coomber *et al*., 2011).

### Pharmacological inactivation of POm in vivo

In some experiments, we silenced the activity of POm by pressure injection of lidocaine. A glass patch pipette (tip diameter 20 μm) was back-filled with mineral oil and then front-loaded with 10% lidocaine in ACSF. These experiments required craniotomies over SC, POm and vS1. Extracellular recordings from vS1 neurons were performed using 4-channel linear Neuronexus silicon electrodes inserted at an oblique angle of 45° in vS1, while SC activation was achieved using an optic fiber inserted vertically into the SC. The pipette containing lidocaine was then inserted into POm. A vibrating mesh contacted most of the whiskers on the contralateral side. Control data was collected without any pressure applied to the back end of the lidocaine-containing pipette in POm. Subsequently, 100-200 nl of 10% lidocaine was injected into POm using a Nanoject to inactivate POm while recording from the same neurons in vS1.

### Brain slice recordings

Three to four weeks after viral injection of AAV1-hSyn.ChR2(H134R)-eYFP.WPRE.hGH into SC or AAV1-Ef1a-DIO-hChR2(E123A)-EYFP into POm, mice were deeply anesthetized with isoflurane (3% in oxygen) and immediately decapitated. The brain was quickly extracted and sectioned in a chilled cutting solution containing (in mM): 100 Choline Chloride, 11.60 N-ascorbate, 7 MgCl_2_, 3.10 Na-pyruvate, 2.50 NaH_2_PO_4_ and 0.50 CaCl_2_ (pH = 7.4). Coronal slices at 300 μm thickness containing either SC, POm or vS1 were prepared using a Leica Vibratome 1000S. Slices were incubated in an incubating solution containing (in mM): 92 NaCl, 2.5 KCl, 1.2 NaH_2_PO_4_, 30 NaHCO_3_, 3 sodium pyruvate, 2 CaCl_2_, 2 MgSO_4_ and 25 glucose at 35°C for 30 min, followed by incubation at room temperature for at least 30 minutes before recording. All solutions were continuously bubbled with 95% O2/5% CO2 (Carbogen).

Whole-cell patch-clamp recordings were made under visual control from SC, POm, VPM or vS1 neurons using infrared-differential interference contrast optics (Stuart *et al*., 1993; Landisman & Connors, 2007). During recording, slices were constantly perfused at ~2 ml/min with carbogen-bubbled artificial cerebral spinal fluid (ACSF) containing (in mM): 125 NaCl, 25 NaHCO_3_, 3 KCl, 1.25 NaH_2_PO_4_, 2 CaCl_2_, 1 MgCl_2_ and 25 glucose maintained at 30-34°C. Patch pipettes were pulled from borosilicate glass and had open tip resistances of 5-7 MΩ when filled with an internal solution containing (in mM): 130 K-Gluconate, 10 KCl, 10 HEPES, 4 MgATP, 0.3 Na2GTP, 10 Na_2_Phosphocreatine and 0.3% biocytin (pH 7.25 with KOH). All recordings were made in current-clamp using a BVC-700A amplifier (Dagan Instruments, USA). Data were filtered at 10 kHz and acquired at 50 kHz by a Macintosh computer running Axograph X acquisition software (Axograph Scientific, Sydney, Australia) using an ITC-18 interface (Instrutech/HEKA, Germany).

Hyperpolarizing and depolarizing current steps (−200 pA to +600 pA; intervals of 50 pA) were applied via the somatic recording pipette to characterize passive and active properties of neurons. Brain slices were continuously bathed in Gabazine (10 μM) to block inhibition mediated by GABA_A_ receptors. Other pharmacological agents used in these experiments included tetrodotoxin (TTX; 1μM) and 4-aminopyridine (4-AP; 100μM), as noted in the Results. For photo-stimulation of ChR2-expressing neurons and axon terminals a 470 nm LED (ThorLabs) was mounted on the epi-fluorescent port of the microscope (Olympus BX50) allowing wide-field illumination through the microscope objective. The timing, duration and strength of LED illumination was controlled by the data acquisition software (Axograph).

### Data analysis and statistics

Data analysis was performed using custom programs in MATLAB (Mathworks, Natick, MA) or with Axograph X. For *in vivo* recordings the spiking response of each neuron was defined as the number of action potentials within a 100 ms window post stimulus onset (light in SC, whisker defection or both) averaged across 50 repetitions of each stimulus. Peri-stimulus time histograms (PSTH; 1 ms bin width) were constructed for the different stimulus conditions. Response latency was defined as the first occurrence, after stimulus onset, of two consecutive bins in the PSTH with significant responses (t-test; p < 0.05). The background, spontaneous firing rate of each neuron was calculated in a 150 ms interval before the stimulus onset. To determine if a neuron responded to a stimulus we used nonparametric, receiver operating characteristic (ROC) analysis (Green & Swets, 1966). Formally, ROC estimates how well an ideal observer can classify whether a given spike count was recorded in one of two possible conditions: Here, the absence or presence of light /whisker stimulation (or both). In each case we compared the trial-by-trial spike count after stimulation onset with that observed prior to stimulation onset. The overlap between these two spike count distributions was quantified by applying criterion levels ranging from the minimum to the maximum observed spike count. The statistical significance of the ROC value was determined by bootstrap analysis, with ROC calculated 1000 times using the same experimental data randomly assigned to the experimental condition. The fraction of bootstrapped ROC values greater than the observed value indicates the level of significance. ROC analysis was used to determine the responsivity of vS1 neurons to whisker stimulation and/or SC activation. The impact of SC activation on the whisker input-output relationship of vS1 neurons was only quantified for neurons that had a statistically significant response to at least one of the whisker intensities tested and if their response to whisker stimulation was statistically significantly enhanced when SC was activated optogenetically by light.

For paired data Wilcoxon’s non-parametric matched pairs test or a paired t-test were used to test statistical significance. Statistical significance was set at p < 0.05. Results are presented as average values ± the standard error of the mean (SEM), unless otherwise stated. In the Figures “ns” denotes not statistically significant, whereas an asterisks denotes p < 0.05.

## Acknowledgments

We would like to thank Randy Bruno for encouraging us to perform the POm inactivation experiments and for technical advice. This work was supported by the Australian Research Council Centre of Excellence for Integrative Brain Function to E. A. and G.J.S.

## Author Contributions

S.G., E.A. and G.J.S. designed the experiments and interpreted the data. S.G. performed and analyzed the *in vivo* experiments. S.H. performed and analyzed the *in vitro* experiments. S.G. drafted the manuscript and all authors edited and approved the final version of the manuscript.

## References

Ahissar E & Oram T (2015). Thalamic Relay or Cortico-Thalamic Processing? Old Question, New Answers. Cereb Cortex 25, 845–848.

Ahmadlou M, Zweifel LS & Heimel JA (2018). Functional modulation of primary visual cortex by the superior colliculus in the mouse. Nat Commun 9, 3895.

Arabzadeh E, Zorzin E & Diamond ME (2005). Neuronal encoding of texture in the whisker sensory pathway. ed. Meister M. PLoS Biol 3, e17.

Barth TM & Schallert T (1987). Somatosensorimotor function of the superior colliculus, somatosensory cortex, and lateral hypothalamus in the rat. Exp Neurol 95, 661–678.

Beltramo R & Scanziani M (2019). A collicular visual cortex: Neocortical space for an ancient midbrain visual structure. Science (80-) 363, 64 LP–69.

Berman RA & Wurtz RH (2011). Signals Conveyed in the Pulvinar Pathway from Superior Colliculus to Cortical Area MT. J Neurosci 31, 373–384.

Bosman LWJ, Houweling AR, Owens CB, Tanke N, Shevchouk OT, Rahmati N, Teunissen WHT, Ju C, Gong W, Koekkoek SKE & De Zeeuw CI (2011). Anatomical pathways involved in generating and sensing rhythmic whisker movements. Front Integr Neurosci 5, 53.

Brecht M (2007). Barrel cortex and whisker-mediated behaviors. Curr Opin Neurobiol 17, 408–416.

Castejon C, Barros-Zulaica N, Nuñez A, Rudy B, Fishell G & Wu C (2016). Control of Somatosensory Cortical Processing by Thalamic Posterior Medial Nucleus: A New Role of Thalamus in Cortical Function. PLoS One 11, e0148169.

Castro-Alamancos M & Keller A (2011). Vibrissal midbrain loops. Scholarpedia 6, 7274.

Castro-Alamancos MA & Favero M (2016). Whisker-related afferents in superior colliculus. J Neurophysiol 115, 2265–2279.

Cavanaugh J, Alvarez BD & Wurtz RH (2006). Enhanced Performance with Brain Stimulation: Attentional Shift or Visual Cue? J Neurosci 26, 11347–11358.

Cohen JD, Hirata A & Castro-Alamancos M a (2008). Vibrissa sensation in superior colliculus: wide-field sensitivity and state-dependent cortical feedback. J Neurosci 28, 11205–11220.

Coomber B, Edwards D, Jones SJ, Shackleton TM, Goldschmidt J, Wallace MN & Palmer AR (2011). Cortical inactivation by cooling in small animals. Front Syst Neurosci 5, 53.

Deschênes M, Veinante P & Zhang Z-W (1998). The organization of corticothalamic projections: reciprocity versus parity. Brain Res Rev 28, 286–308.

Diamond ME & Arabzadeh E (2013). Whisker sensory system - From receptor to decision. Prog Neurobiol 103, 28–40.

Dräger UC & Hubel DH (1976). Topography of visual and somatosensory projections to mouse superior colliculus. J Neurophysiol 39, 91–101.

Furuta T, Kaneko T & Deschênes M (2009). Septal neurons in barrel cortex derive their receptive field input from the lemniscal pathway. J Neurosci 29, 4089–4095.

Gambino F, Pagès S, Kehayas V, Baptista D, Tatti R, Carleton A & Holtmaat A (2014). Sensory-evoked LTP driven by dendritic plateau potentials in vivo. Nature 515, 116–119.

Gharaei S, Arabzadeh E & Solomon SG (2018). Integration of visual and whisker signals in rat superior colliculus. Sci Rep 8, 16445.

Green DM & Swets JA (1966). Signal detection theory and psychophysics. Wiley New York.

Heaton JT, Sheu SH, Hohman MH, Knox CJ, Weinberg JS, Kleiss IJ & Hadlock TA (2014). Rat whisker movement after facial nerve lesion: evidence for autonomic contraction of skeletal muscle. Neuroscience 265, 9–20.

Hemelt ME & Keller A (2008). Superior colliculus control of vibrissa movements. J Neurophysiol 100, 1245–1254.

Herman JP & Krauzlis RJ (2017). Color-Change Detection Activity in the Primate Superior Colliculus. eNeuro; DOI: 10.1523/ENEURO.0046-17.2017.

Kaneshige M, Shibata K, Matsubayashi J, Mitani A & Furuta T (2018). A Descending Circuit Derived From the Superior Colliculus Modulates Vibrissal Movements. Front Neural Circuits 12, 1–12.

Kichula EA & Huntley GW (2008). Developmental and comparative aspects of posterior medial thalamocortical innervation of the barrel cortex in mice and rats. J Comp Neurol 509, 239–258.

Killackey HP & Erzurumlu RS (1981). Trigeminal projections to the superior colliculus of the rat. J Comp Neurol 201, 221–242.

Krauzlis RJ, Lovejoy LP & Zénon A (2013). Superior colliculus and visual spatial attention. Annu Rev Neurosci 36, 165–182.

Landisman CE & Connors BW (2007). VPM and PoM Nuclei of the Rat Somatosensory Thalamus: Intrinsic Neuronal Properties and Corticothalamic Feedback. Cereb Cortex 17, 2853–2865.

Lee CCY, Clifford CWG & Arabzadeh E (2019). Temporal cueing enhances neuronal and behavioral discrimination performance in rat whisker system. J Neurophysiol 121, 1048–1058.

Lee CCY, Diamond ME & Arabzadeh E (2016). Sensory Prioritization in Rats: Behavioral Performance and Neuronal Correlates. J Neurosci 36, 3243–3253.

Lomber SG, Payne BR & Horel JA (1999). The cryoloop: an adaptable reversible cooling deactivation method for behavioral or electrophysiological assessment of neural function. J Neurosci Methods 86, 179–194.

Lovejoy LP & Krauzlis RJ (2010). Inactivation of primate superior colliculus impairs covert selection of signals for perceptual judgments. Nat Neurosci 13, 261–266.

Margrie T, Brecht M & Sakmann B (2002). In vivo, low-resistance, whole-cell recordings from neurons in the anaesthetized and awake mammalian brain. Pfl□gers Arch Eur J Physiol 444, 491–498.

May PJ (2006). The mammalian superior colliculus: laminar structure and connections. Prog Brain Res 151, 321–378.

McBride EG, Lee S-YJ & Callaway EM (2019). Local and Global Influences of Visual Spatial Selection and Locomotion in Mouse Primary Visual Cortex. Curr Biol 29, 1592–1605.e5.

McHaffie JG & Stein BE (1982). Eye movements evoked by electrical stimulation in the superior colliculus of rats and hamsters. Brain Res 247, 243–253.

Mease RA, Metz M & Groh A (2016). Cortical Sensory Responses Are Enhanced by the Higher-Order Thalamus. Cell Rep 14, 208–215.

Muller JR, Philiastides MG & Newsome WT (2005). Microstimulation of the superior colliculus focuses attention without moving the eyes. Proc Natl Acad Sci U S A 102, 524–529.

Ohno S, Kuramoto E, Furuta T, Hioki H, Tanaka YR, Fujiyama F, Sonomura T, Uemura M, Sugiyama K & Kaneko T (2012). A Morphological Analysis of Thalamocortical Axon Fibers of Rat Posterior Thalamic Nuclei: A Single Neuron Tracing Study with Viral Vectors. Cereb Cortex 22, 2840–2857.

Ranjbar-Slamloo Y & Arabzadeh E (2017). High-velocity stimulation evokes “dense” population response in layer 2/3 vibrissal cortex. J Neurophysiol 117, 1218–1228.

Robinson DL & Petersen SE (1992). The pulvinar and visual salience. Trends Neurosci 15, 127–132.

Rodman HR, Gross CG & Albright TD (1990). Afferent basis of visual response properties in area MT of the macaque. II. Effects of superior colliculus removal. J Neurosci 10, 1154.

Roger M & Cadusseau J (1984). Afferent connections of the nucleus posterior thalami in the rat, with some evolutionary and functional considerations. J Hirnforsch 25, 473–485.

Rowland BA, Quessy S, Stanford TR & Stein BE (2007). Multisensory integration shortens physiological response latencies. J Neurosci 27, 5879–5884.

Sabri MM & Arabzadeh E (2018). Information Processing Across Behavioral States: Modes of Operation and Population Dynamics in Rodent Sensory Cortex. Neuroscience 368, 214–228.

Sachidhanandam S, Sreenivasan V, Kyriakatos A, Kremer Y & Petersen CCH (2013). Membrane potential correlates of sensory perception in mouse barrel cortex. Nat Neurosci 16, 1671–1677.

Semba K & Egger MD (1986). The facial ?motor? nerve of the rat: Control of vibrissal movement and examination of motor and sensory components. J Comp Neurol 247, 144–158.

Smith DC & Spear PD (1979). Effects of superior colliculus removal on receptive-field properties of neurons in lateral suprasylvian visual area of the cat. J Neurophysiol.

Snow JC, Allen HA, Rafal RD & Humphreys GW (2009). Impaired attentional selection following lesions to human pulvinar: evidence for homology between human and monkey. Proc Natl Acad Sci U S A 106, 4054–4059.

Sparks DL (1999). Conceptual issues related to the role of the superior colliculus in the control of gaze. Curr Opin Neurobiol 9, 698–707.

Stein BE (2012). The new handbook of multisensory processes. The MIT Press.

Stepniewska I, Qi H-X & Kaas JH (1999). Do superior colliculus projection zones in the inferior pulvinar project to MT in primates? Eur J Neurosci 11, 469–480.

Stuart GJ, Dodt HU & Sakmann B (1993). Patch-clamp recordings from the soma and dendrites of neurons in brain slices using infrared video microscopy. Pflügers Arch Eur J Physiol 423, 511–518.

Tohmi M, Meguro R, Tsukano H, Hishida R & Shibuki K (2014). The Extrageniculate Visual Pathway Generates Distinct Response Properties in the Higher Visual Areas of Mice. Curr Biol 24, 587–597.

Towal RB & Hartmann MJ (2006). Right-Left Asymmetries in the Whisking Behavior of Rats Anticipate Head Movements. J Neurosci 26, 8838–8846.

Triplett JW, Phan A, Yamada J & Feldheim DA (2012). Alignment of multimodal sensory input in the superior colliculus through a gradient-matching mechanism. J Neurosci 32, 5264–5271.

Viaene AN, Petrof I & Murray Sherman S (2011). Properties of the thalamic projection from the posterior medial nucleus to primary and secondary somatosensory cortices in the mouse. PNAS; DOI: 10.1073/pnas.1114828108.

Welker E, Hoogland P V & Van der Loos H (1988). Organization of feedback and feedforward projections of the barrel cortex: a PHA-L study in the mouse. Exp brain Res 73, 411–435.

Wimmer RD, Schmitt LI, Davidson TJ, Nakajima M, Deisseroth K & Halassa MM (2015). Thalamic control of sensory selection in divided attention. Nature 526, 705–709.

Wise SP & Jones EG (1977). Somatotopic and columnar organization in the corticotectal projection of the rat somatic sensory cortex. Brain Res 133, 223–235.

Zénon A & Krauzlis RJ (2012). Attention deficits without cortical neuronal deficits. Nature 489, 434–437.

Zhang W & Bruno RM (2019). High-order thalamic inputs to primary somatosensory cortex are stronger and longer lasting than cortical inputs ed. Calabrese RL. Elife 8, e44158.

Zingg B, Chou X, Zhang Z, Mesik L, Liang F, Tao HW & Zhang LI (2017). AAV-Mediated Anterograde Transsynaptic Tagging: Mapping Corticocollicular Input-Defined Neural Pathways for Defense Behaviors. Neuron 93, 33–47.

