## Supplementary materials for "Superior colliculus modulates cortical coding of somatosensory information"

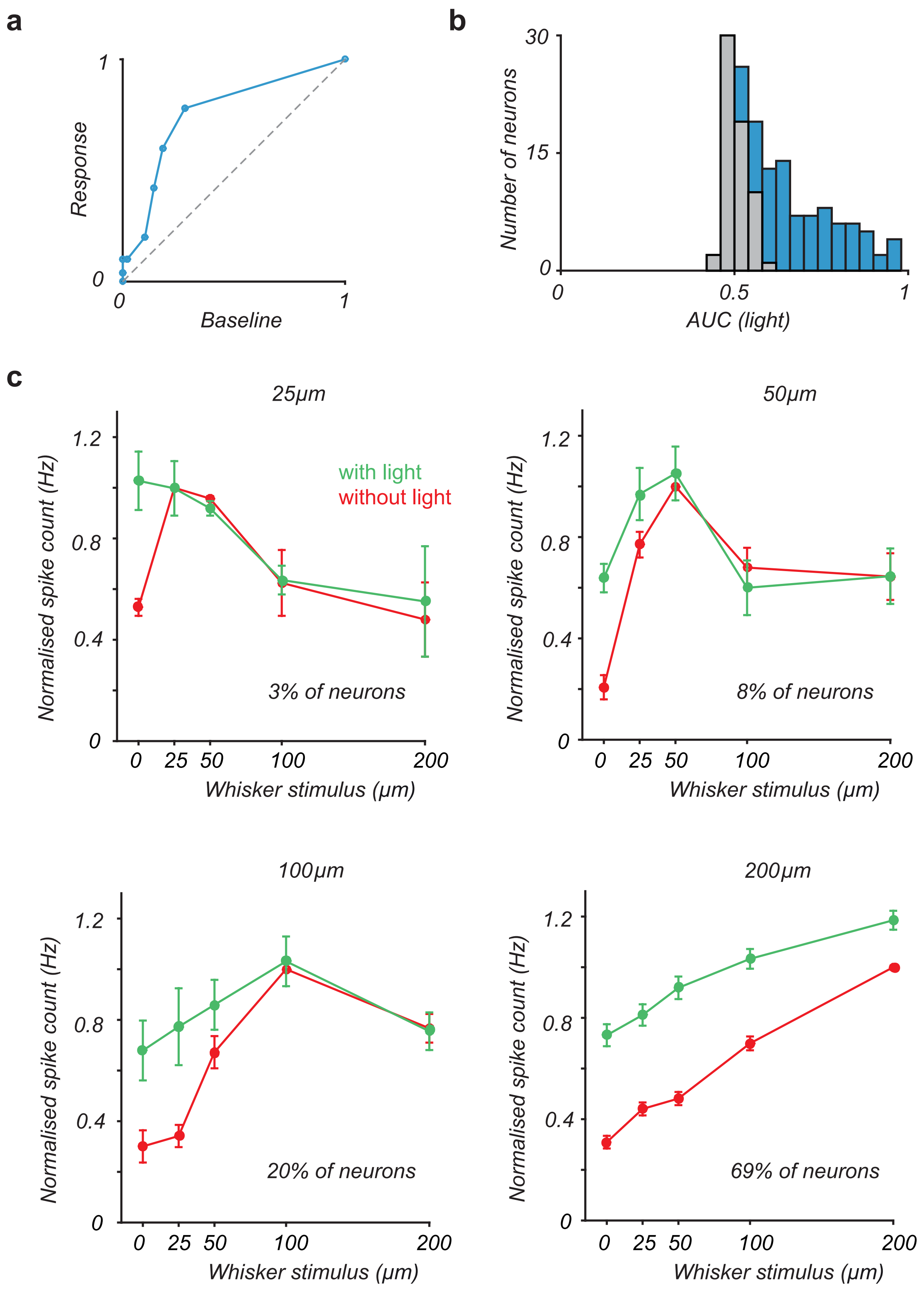


Supplementary Figure 1

**Supplementary Fig. 1: ROC analysis and impact on SC on whisker tuning. a.** ROC curve for an example vS1 neuron during SC light stimulation. The dashed line shows what is expected by chance. For this neuron, the area under the ROC curve is 0.76. **b.** The distribution of areas under the ROC curve across all neurons (n=149) where gray depicts vS1 neurons where there was no significant change in spiking and blue (n=87) shows vS1 neurons with a significant increase in spiking in response to SC stimulation. **c.** Whisker responsive vS1 neurons that also responded to SC activation (n=101) were divided into 4 groups depending on the whisker stimulus that evoked the maximum response (either 25, 50, 100 and 200 μm). Spiking activity of each group was normalized to the maximum response to the whisker stimulation alone. Error bars represent SEM. Seventy neurons (69%) had the greatest response to 200 μm whisker deflections (bottom right), 20 neurons (20%) had the greatest response to 100 μm whisker deflections (Bottom, left), 8 neurons (8%) had the greatest response to 50 μm whisker defections (top, right) and 3 neurons (3%) had the greatest response to 25 μm defections (top, left). For every group, optogenetic activation of SC only impacted on whisker responses for deflection amplitudes lower than that which evoked the maximum response.

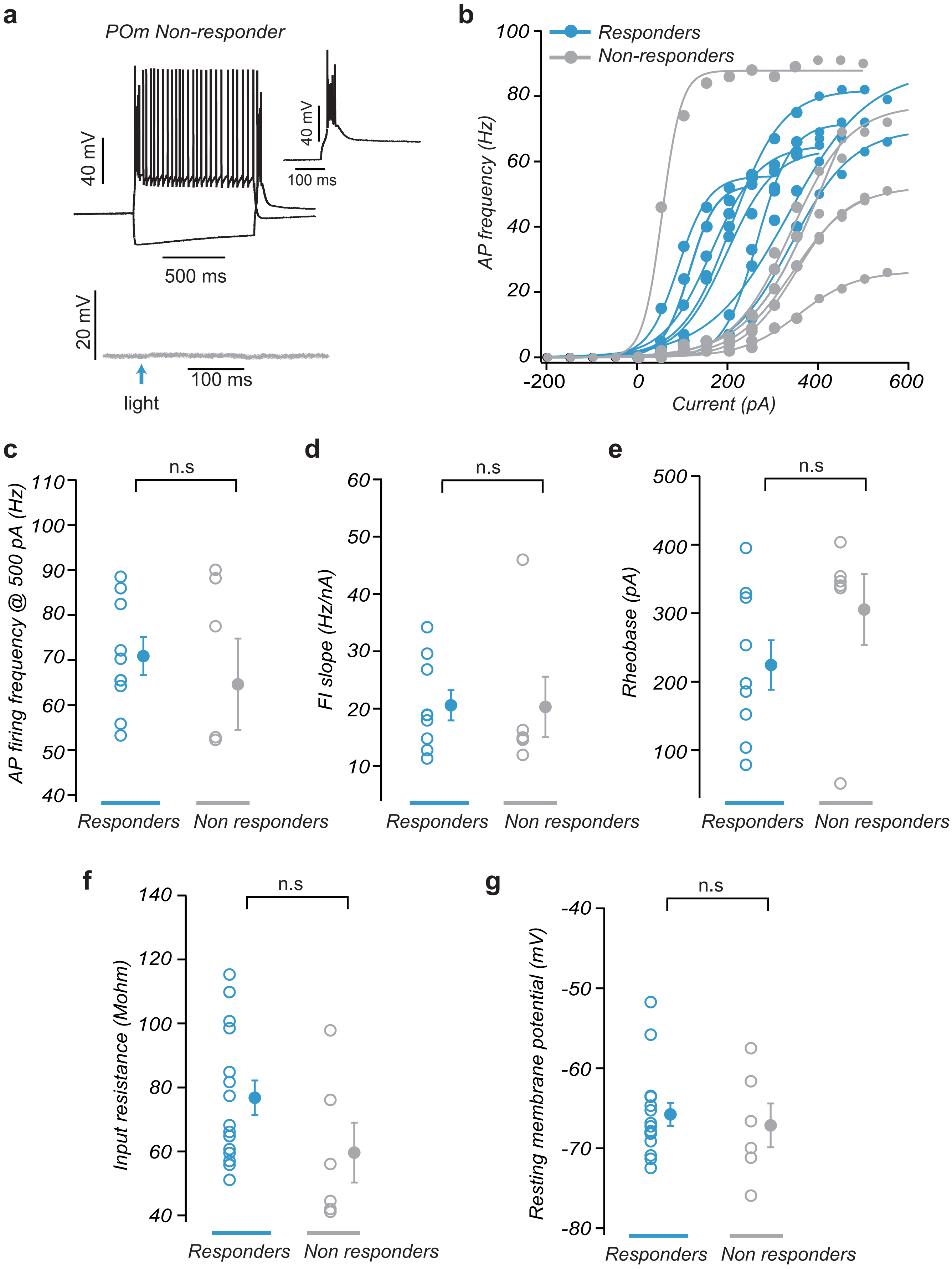


Supplementary Figure 2

**Supplementary Fig. 2: POm neurons that receive inputs from SC have similar active and passive properties to those that do not. a.** Top: Response of a POm neuron to somatic depolarizing (+300 pA) and hyperpolarizing (-400 pA) current steps. Inset shows the rebound spikes. Bottom: This POm neuron does not receive SC input. **b.** Plot of action potential (AP) firing frequency versus somatic current (f/I) in POm neurons that receive direct SC input (“responders”; blue; n=16) and those that do not (“non-responders”; grey; n=6). **c.** Comparison of AP firing frequency in response to a +500 pA somatic current injection in “responders” and “non-responders”. **d.** Comparison of f/I slope in “responders” and “non-responders” shown in panel B. **e.** Comparison of the minimum current required to evoke APs (rheobase) in “responders” and “non-responders”. **f,g.** Comparison of input resistance **(f)** and resting membrane potential **(g)** in “responders” and “non-responders”. Pooled data represents mean ± SEM.

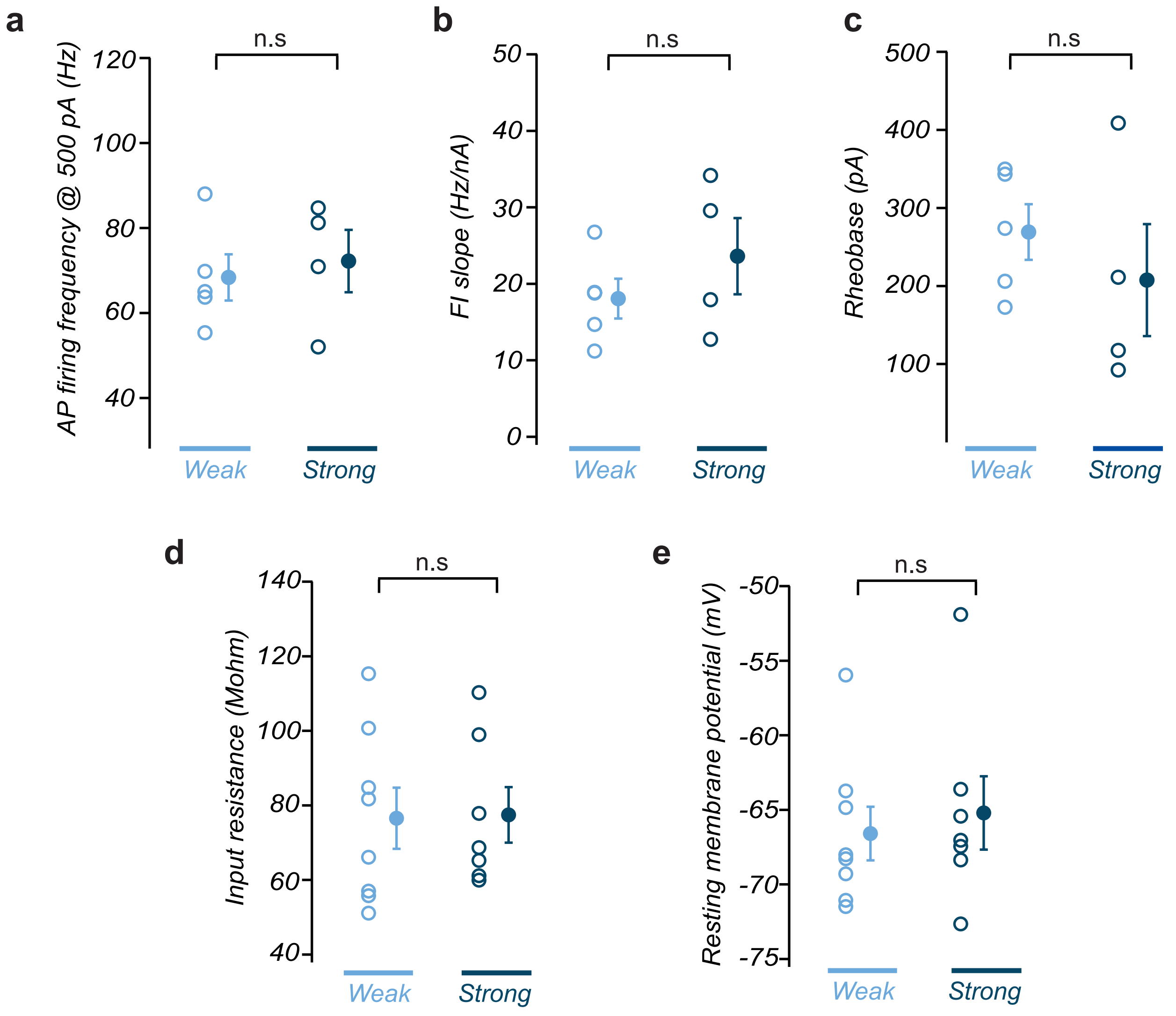


Supplementary Figure 3

**Supplementary Fig. 3: POm neurons that receive weak and strong SC input have similar active and passive properties. a.** Comparison of AP firing frequency in response to a +500 pA somatic current injection in POm neurons receiving weak and strong SC input. **b.** Comparison of f/I slope in POm neurons receiving weak and strong SC input. **c.** Comparison of the minimum current required to evoke APs (rheobase) in POm neurons receiving weak and strong SC input. **d,e.** Comparison of input resistance **(d)** and resting membrane potential **(e)** in POm neurons receiving weak and strong SC input. Pooled data represents mean ± SEM.
